# Genomic factors shaping codon usage across the Saccharomycotina subphylum

**DOI:** 10.1101/2024.05.23.595506

**Authors:** Bryan Zavala, Lauren Dineen, Kaitlin J. Fisher, Dana A. Opulente, Marie-Claire Harrison, John F. Wolters, Xing-Xing Shen, Xiaofan Zhou, Marizeth Groenewald, Chris Todd Hittinger, Antonis Rokas, Abigail Leavitt LaBella

## Abstract

Codon usage bias, or the unequal use of synonymous codons, is observed across genes, genomes, and between species. The biased use of synonymous codons has been implicated in many cellular functions, such as translation dynamics and transcript stability, but can also be shaped by neutral forces. The Saccharomycotina, the fungal subphylum containing the yeasts *Saccharomyces cerevisiae* and *Candida albicans*, has been a model system for studying codon usage. We characterized codon usage across 1,154 strains from 1,051 species to gain insight into the biases, molecular mechanisms, evolution, and genomic features contributing to codon usage patterns across the subphylum. We found evidence of a general preference for A/T-ending codons and correlations between codon usage bias, GC content, and tRNA-ome size. Codon usage bias is also distinct between the 12 orders within the subphylum to such a degree that yeasts can be classified into orders with an accuracy greater than 90% using a machine learning algorithm trained on codon usage. We also characterized the degree to which codon usage bias is impacted by translational selection. Interestingly, the degree of translational selection was influenced by a combination of genome features and assembly metrics that included the number of coding sequences, BUSCO count, and genome length. Our analysis also revealed an extreme bias in codon usage in the Saccharomycodales associated with a lack of predicted arginine tRNAs. The order contains 24 species, and 23 are computationally predicted to lack tRNAs that decode CGN codons, leaving only the AGN codons to encode arginine. Analysis of Saccharomycodales gene expression, tRNA sequences, and codon evolution suggests that extreme avoidance of the CGN codons is associated with a decline in arginine tRNA function. Codon usage bias within the Saccharomycotina is generally consistent with previous investigations in fungi, which show a role for both genomic features and GC bias in shaping codon usage. However, we find cases of extreme codon usage preference and avoidance along yeast lineages, suggesting additional forces may be shaping the evolution of specific codons.

## Introduction

The genetic code is degenerate, and all but two amino acids are encoded by multiple synonymous codons. It is consistently observed that the use of synonymous codons is biased within genes, between genes within a genome, and between species (*1, 2*). The unequal use of synonymous codons is known as codon usage bias (CUB) and has been observed widely across organisms (*1*). Some of these biases are associated with the functional properties of codons. Generally, the relative frequency of synonymous codons is proportional to the number of available tRNAs with a corresponding anticodon, which facilitates translation (*3*). Transcripts containing codons decoded by abundant tRNAs are also frequently expressed at a higher level (*4*). Codon usage bias has been found to have a functional role in many cellular processes, including mRNA stability (*5*), transcriptional control (*6*), protein folding (*7*), chromatin availability (*8*), and ribosome dynamics (*9*).

The involvement of codon usage bias in diverse cellular processes suggests that codon usage is under natural selection. Natural selection acting on codon usage is typically attributed to translational selection, a form of adaptation associated with increased translational efficiency in highly expressed genes accomplished by tuning CUB to the most abundant tRNAs (*10*). Non-adaptive forces such as GC-biased gene conversion, mutational bias, and genetic drift also have a signature effect on the nucleotide composition landscape. GC-biased gene conversion is a process that generally influences eukaryotes and is due to recombination that prefers the transmission of GC alleles, thus increasing the GC content (*11*). However, biased gene conversion may not significantly impact GC content in some species, such as *Saccharomyces cerevisiae* (*12*). Mutational biases can also impact global GC composition. Mutational biases are driven by differential transition and transversion rates (*13*), mutational pressure towards AT content driven by the deamination of cytosine to uracil (*14*), and replication strand bias (*15*). Finally, genetic drift decreases selection efficiency, including translational selection on codon usage (*16*).

The relative degree to which adaptive and non-adaptive forces shape codon usage bias differs across the tree of life. In some species, especially those with small population sizes, neutral forces are thought to predominately shape codon usage biases (*16*). In this case, codon usage is highly correlated with global GC content and is not predictive of gene expression (*10*). Other species, however, exhibit a strong signature of translational selection such that codon composition is highly predictive of gene expression level (*17*). Methods have been developed to disentangle these forces to assess the relative contribution of translational selection on genomic codon usage bias (*10, 18, 19*). These methods have found a wide range of translational selection levels across organisms (*10*). This variation has been attributed to factors such as genome size, total number of genomic tRNAs (*10*), and effective population size (*16*). These features, however, do not fully explain the observed variation, as numerous exceptions exist (*20*). It remains to be seen if other metabolic, phenotypic, or genomic traits impact the degree of translational selection on CUB or CUB itself.

The fungal subphylum Saccharomycotina, one of three subphyla in the phylum Ascomycota, is an excellent model system for studying codon usage evolution (*20–22*). Codon usage in the Saccharomycotina is among the most unusual in eukaryotes as it harbors three orders (Serinales, Alaninales, Alloscoideales) that have undergone nuclear codon reassignments in which the CTG codon encodes for serine or alanine, instead of leucine (*23–25*). The extensive genomic diversity of these yeasts provides insight into the evolution of codon usage and possible factors shaping them. Previous research on a subset of the subphylum of Saccharomycotina (332 or fewer yeasts) has revealed that most, but not all, species within this group are subject to high levels of translational selection (*20, 26, 27*). This work found that genomic tRNA copy number was the most robustly associated with levels of translational selection on codon usage, with the highest levels occurring at intermediate tRNA levels (*20*).

Surprisingly, genome size was not highly correlated with levels of translational selection. There are likely other forces shaping codon usage bias within this diverse subphylum. These findings warrant additional investigation in the entire yeast subphylum, which the Y1000+ Project has recently made possible through the generation of genomic and phenotypic characterization of 1,154 yeast strains from 1,051 species—nearly all known species of yeasts (*28*).

To understand the evolution of codon usage bias in Saccharomycotina, we analyzed the genomic-wide codon usage metrics and genomic features across the subphylum. We measured relative synonymous codon usage (RSCU), which reflects codon preference, and the association between RSCU and various genomic features. Across the subphylum, we found a general preference towards AT-ending codons, but there were significant differences in codon usage between species. A phylogenetic principal component analysis (pPCA) revealed distinct patterns of RSCU values that differentiated the Serinales and Dipodascales from the rest of the subphylum. A random forest classifier was able to classify strains into their taxonomic orders with high accuracy based solely on RSCU values, indicating the presence of distinct patterns in codon usage patterns in the subphylum. Further phylogenetic statistical analysis on codon usage resulted in significant correlations with GC content metrics but also revealed intriguing correlations with specific genomic features. Significant correlations were found between translational selection levels (measured by S-Value) and tRNA count, assembly metrics, and the number of protein-coding sequences. We also conducted an in-depth analysis of RSCU values of CGN codons, which revealed that no tRNA was computationally predicted to decode CGN codons in the Saccharomycodales. The Saccharomycodales contained 23 species (out of 24) that were predicted to lack CGN decoding tRNA genes. We conducted additional analyses on gene expression, tRNA alignments, and conserved arginine sites, further suggesting the loss of functional CGN codons. This analysis of codon usage throughout the subphylum highlights the diversity of codon usage strategies and identifies some of the genomic features that may constrain this diversity.

## Materials & Methods

### Codon usage data and metrics

The protein-coding sequence annotations of 1,154 yeast strains from 1,051 species were collected from a previous study from the Y1000+ Project (http://y1000plus.org) of the yeast genomes in the subphylum Saccharomycotina (Table S1)(*28*). Codon calculations were generated through the sequence analysis tool EMBOSS v6.6.0.0 (*29*), which calculated codon frequencies, percentages, and counts for every coding sequence in each yeast strain. Most codon analysis software does not include all possible yeast nuclear translation tables and, therefore, is inaccurate for the Saccharomycotina. To address this, the RSCU (Relative Synonymous Codon Usage) for every yeast strain was calculated by in-house scripts (https://github.com/The-Lab-LaBella/RSCU_Calculation_Analysis) that accounted for the nuclear codon reassignments in the Serinales, Alaninales, and Ascoideales. The RSCU is the observed number of occurrences of a synonymous codon divided by the expected number of occurrences if codon usage was random (*4*).The RSCU values for every protein-coding sequence were used to calculate each genome’s genome-wide average RSCU values.

To estimate the level of translational selection acting on codon usage in our yeast genomes, we applied the S-test to calculate the S-value proposed by dos Reis et al. (*10*) using the tAI package in R (*10*). We used this package to generate tRNA adaptation index (tAI) values for every gene. The S-value for each genome was then calculated as the correlation between tAI and a combination of the synonymous third codon position GC content (GC3s) and the effective number of codons. This method was chosen because we could conduct high-throughput analysis on the command line instead of a graphical user interface. Other methods have shown similar trends in detecting selection on codon usage in the yeasts (*18*).

### tRNA Analysis

tRNAscan-SE 2.0.9 (*30*) was used to predict tRNAs in the genome for each strain using the standard eukaryotic parameters. We generated a filtered tRNAscan-SE dataset in which we removed pseudogenes, tRNAs lacking isotypes, and tRNAs containing mismatches between anticodon and predicted isotype. Serine or alanine tRNAs with the CAG anticodon were included for the Serinales, Alaninales, and Ascoideales as the orders have undergone a reassignment of the canonical codon (CUG) for leucine to serine (Serinales (*31*) and Ascoideales (*32*)) or alanine (Alaninales (*23, 24*)). After filtering, the total tRNA-ome size was calculated as the sum of all tRNAs. We conducted additional analyses on the tRNAs of Saccharomycodales yeasts, specifically analyses of the arginine-decoding tRNAs. First, we analyzed the mitochondrial tRNA content of several yeast species in the Saccharomycodales. The mitochondrial genomes were obtained from a recent study (*33*) and they were analyzed for tRNA content using tRNA-scan with the organelle option.

The conservation of Saccharomycodales arginine tRNAs was also assessed using the tRNAviz web browser (*34*) . In addition to visualization, tRNAviz calculates a penalty score for each position in the supplied tRNAs. Scores range from 0 (identical to the reference) to -15, indicating a highly divergent site. The reference set used in this analysis was the pre-computed Saccharomycotina dataset.

Annotations of tRNA modification enzymes were obtained from KEGG (*35*) annotations previously conducted (*28*)The annotations examined were KEGG Ortholog groups K15440 (TAD1), K15441 (TAD2), and K15442 (TAD3). They were examined manually to identify any annotation errors.

### Genomic and Phenotypic Traits of Yeasts

Next, we aimed to identify genomic and phenotypic traits that may influence levels of codon usage bias and translational selection in yeasts. To do this, we used a variety of yeast traits obtained from the publication of the genomes (*28*). Genomic traits analyzed were GC content metrics, tRNA-ome size, S-value, genome size, genome assembly metrics, BUSCO completeness metrics, and number of protein-coding sequences. Phenotypic traits analyzed were metabolic niche breadth(*28*). Metabolic niche breadth is the total number of carbon or nitrogen substrates on which a yeast strain can grow.

### Visualization and Phylogenetic Statistical Analyses

The genome-wide average RSCU values across the subphylum were analyzed with hierarchical clustering performed using the ComplexHeatmap package (*36*). To decipher patterns and covariance across strains in their codon usage variation, a phylogenetic principal component (pPCA) analysis was performed in R using the phytools package (*37*). A pPCA was used to take into account the non-independence of closely related species. The phylogeny of all 1,154 yeasts, which was constructed using maximum likelihood analysis of a concatenated alignment of 1,403 single-copy orthologs, was obtained from the previous study (*28*). Similarly, phylogenetic generalized least squares (PGLS) and phylogenetic independent contrasts (PIC) were used to estimate correlations between biological features, genomic features, and RSCU. The RSCU versus RSCU analysis was conducted under a PGLS with maximum likelihood estimation of Pagel’s lambda in the caper v1.7.3 package (*38*). In cases where the lambda estimation failed, possibly due to a maximum likelihood outside the bounds of 1×10^-6^ and 1, it was set to 1 to ensure the most conservative analysis. The yeast feature versus yeast feature analysis was also conducted using the same PGLS method. In the feature analysis, two genomes were dropped due to naming issues: genome names yHMPu5000026270_Candida_sp._SPAdes, yHMPu5000034970_Candida_sp._SPAdes. Due to differences in scale, the feature versus RSCU analysis was conducted using a PIC in the ape v5.7-1 package (*39*). Strains without metabolic breadth measurements were removed before phylogenetic comparative analysis.

### RSCU Random Forest Classifier

To determine if yeast strains can be classified by codon usage, a random forest classifier model was created to determine the order of a particular yeast strain based solely on their genome-wide average RSCU values. The model was built from the R package randomForest v.4.7-1.1 (*40*) with a matrix comprising the genome-wide average RSCU values from 59 codons for each strain. The model was trained with 70% of the data, and 30% was withheld for testing. The trained model included information regarding the significant variables (codons) in classifying yeast strains to orders and error rate stabilization.

### Gene Expression Analysis

We conducted mRNA sequencing to verify the presence of rare codons in the expressed genes of Saccharomycodales. Triplicate cultures *of Hanseniaspora occidentals* var. *occidentals* and *Hanseniaspora uvarum* were grown for 18 hours in 20 ml rich YPD medium (yeast extract, peptone, 2% glucose) in a room-temperature shaker at 250 rpm. Cell pellets were flash-frozen and stored at -80°C. mRNA was isolated using the NEBNext Poly(A) mRNA Magnetic Isolation Module (NEB) with the NEBNext Ultra II Directional RNA Library Prep kit (NEB). One of the three *H. occidentals* var. *occidentals* libraries failed, leaving only two replicates for sequencing. Sequencing was performed at the University of Wisconsin-Madison Biotechnology Center.

Paired-end reads were trimmed and deduplicated using fastp v.0.20.1 (*41*). A guided assembly was conducted using HISAT2 v2.2.1 (*42*) to align the sequence reads against their respective reference draft genomes, both of which were retrieved from the Y1000+ Project (*28*). The resulting alignment files were processed through Stringtie v2.2.1 (*43*) for transcript annotation from their respective reference species annotation files, followed by extracting the protein-coding sequences from the reference genomes of each replicate annotation file into FASTA files using TransDecoder v5.5.0. Putative transcript functions were assigned using the NCBI Blast web search (*44*).

### Conserved Arginine Site Analysis

Our analyses revealed an extreme bias in arginine codons in the Saccharomycodales.

Therefore, we explored the evolutionary context of conserved arginine positions within the genomes. We identified highly conserved arginine amino acid positions in the 1,403 single-copy orthologs previously used to build the Saccharomycotina phylogeny (*28*). The DNA sequences of these orthologs were translated into amino acid sequences using EMBOSS *transeq* (*29*) and then aligned using mafft v7.273 (*45*) with the “fftnsi ’’ settings. The DNA coding sequences were then aligned by codon using the protein alignment as a reference using a custom script available on the Figshare repository. Conserved arginine positions, defined by a simple majority of sequences, were identified in the amino acid alignments using the EMBOSS *cons* function.

We then focused on the Saccharomycodales, the Saccharomycodales (the sister order to the Saccharomycodales), and the Phaffomycetales (an outgroup.) We identified sites in the alignment where 80% or more of the amino acids were arginine, and the exact codon was conserved within each order at 80% or more. This procedure identified highly conserved arginine positions with highly conserved arginine codons. This allowed us to identify patterns of arginine codon usage within this group.

## Results & Discussion

### Codon Usage Variation Across the Subphylum

Our previous work identified significant variation in codon usage in 332 strains of the yeast subphylum (*20*). Here, we have more than tripled the sampling within the subphylum. First, we calculated the strain-level average relative synonymous codon usage (RSCU) values for the 59 degenerate codons. Hierarchical clustering revealed patterns of codon preference throughout the subphylum (Figure 1, Table S2, Figure S1). We observed a general preference (RSCU > 1) for A/T-ending codons, while G/C-ending codons were unpreferred (RSCU < 1). However, there were notable deviations from the general patterns in RSCU values. Specifically, the codon TTG, which codes for leucine, was grouped among the preferred A/T-ending codons and was the only G/C-ending codon generally preferred. Another observation was that the A/T-ending codons of CTT (leucine), CGT (arginine), GTA (valine), ATA (isoleucine), CTA (leucine), and CGA (arginine) were among the unpreferred codons in the subphylum. Other RSCU patterns were specific to one or a few orders of yeasts. The codon AGA (arginine) exhibited an extreme preference throughout all the orders except for Lipomycetales, Trigonopsidales, and Dipodascales. A similar pattern was observed with the TTG codon. The codon TTA was highly variable across the subphylum and exhibited extreme preferences (preferred and unpreferred).

**Figure 1:**
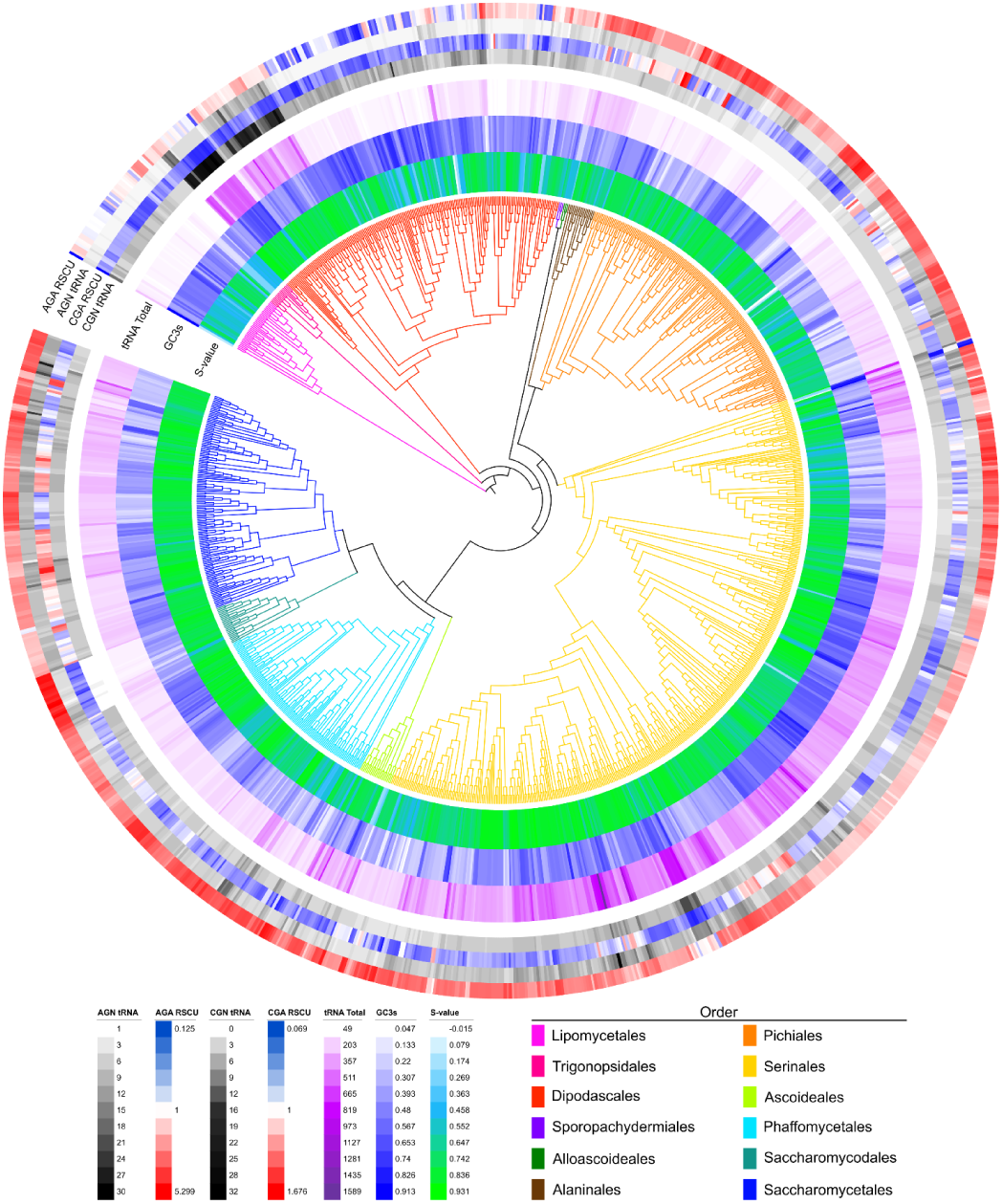
Variation of codon-associated metrics across the Saccharomycotina subphylum. The S-value is a measure of translational selection on codon usage that varies from no selection (0) to high levels of selection (1). The synonymous GC content of the third codon position (GC3s) and total genomic tRNA count are also shown. Representative relative synonymous codon usage (RSCU) and individual tRNA counts for arginine are also shown. The RSCU and counts of decoding tRNAs vary widely across the phylogeny.

We also investigated the correlation between RSCU and other codon-associated traits, including synonymous GC3 (GC3s) composition, translational selection (S-value), and tRNA-ome size (Figure 1, Table S1, Table S2). These traits represent various factors influencing codon usage variation across the subphylum. S-value (a measure of translational selection) varies significantly across the subphylum (Fig 1.) A high level of translational selection on codon usage within a genome is characterized by a high correlation between codon adaptation to genomic tRNA numbers (tAI) and compositional bias. Therefore, a value of 1 indicates a perfect correlation and a high inferred level of translational selection on codon usage. An S-value of approximately 0.5 indicates an intermediate level of translational selection. Across the yeast subphylum, we observed S-values ranging from a low of -0.015 to a high of 0.931 (mean=0.73, median=0.76). GC composition, which can influence codon usage bias in a non-selective way, is also highly variable across yeasts. We observed synonymous GC3 values from a low of 4.7% to a high of 91.3%. The average across the genomes was 42.36% and a median of 43%. The tRNA-ome is another source of codon usage evolution as the repertoire of tRNA copy number is directly implicated in the presence of translation selection (*10*). The median tRNA-ome in the subphylum was 208 tRNAs, ranging from a minimum of 49 tRNAs to a maximum of 1,589 tRNAs. *Candida lidongshanica* and *Aciculoconidium aculeatum* from the Serinales have an unusually large predicted tRNA-ome with 1,589 tRNAs and 1,037 tRNAs, respectively. Further analysis of the number of distinct anticodon types in each strain revealed that only one species, *Martiniozyma abiesophila* in the Pichiales, violates the theoretical minimum of 30 anticodon types (*46*) by containing only 28 anticodon types. The tRNA content analyses present here are limited by the references used to model tRNAs within tRNAscan-SE and potentially incomplete genomic sequences. Additional experimental work is required to fully elucidate the expressed tRNA content across the yeast subphylum.

### RSCU Patterns are a Defining Feature of Yeast Orders

To examine interspecies RSCU codon usage variation, we conducted a phylogenetic principal component analysis (pPCA). The pPCA revealed that most of the variation (63.56%) (Figure 2) between species is driven by G/C- and A/T-ending codons, which is consistent with previous analyses (*20*). The second principal component (11.96%), which differentiates the Serinales/Ascoideales and Dipodascales/Lipomycetales, is driven by a set of codons that include TTG, CTT, CTA, CTG, AGT, TCA, and TCT. In particular, the usage of TTG, CTT, and CTA (all leucine codons) separates and clusters the Serinales and Ascoideales from the rest. This reflects the avoidance of the reassigned CTG codon in these orders. The Dipodascales also exhibited unique clustering driven primarily by codon preferences for CTG (leucine codon), AGT, TCA, and TCT (serine codons). An interesting deviation was seen for *Dipodascopsis tothii* (order Lipomycetales), which is separate from all the other species in the subphylum.

**Figure 2.**
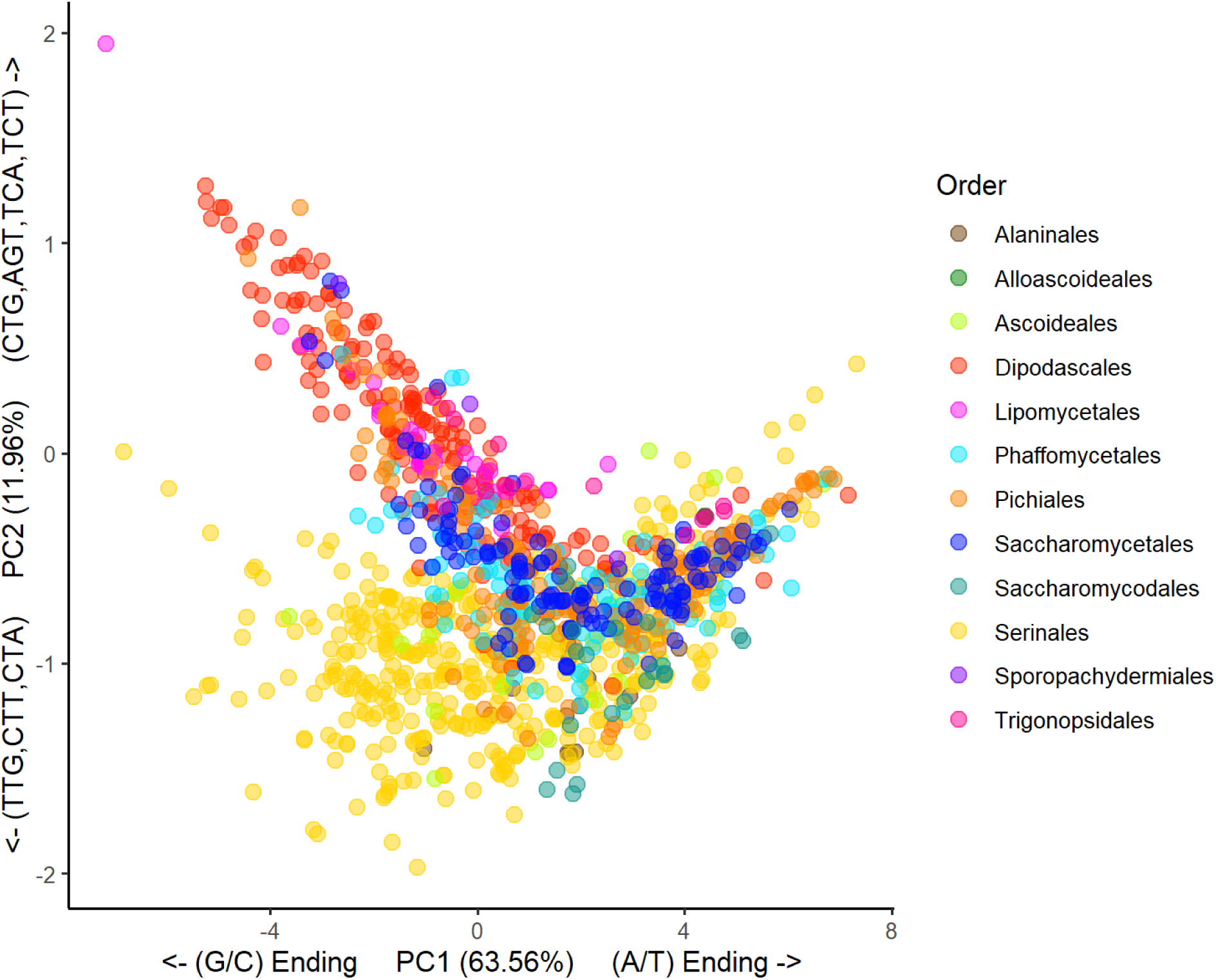
Phylogenetic Principal Component Analysis (pPCA) of the 59 RSCU values of 1,154 yeast strains from 1,051 species. To derive the patterns and covariance of codon usage throughout the subphylum, a pPCA was conducted to determine their relationships. A pPCA was used to take into account the non-independence of biological traits due to phylogeny. The results demonstrated that PC1, which explains 63.56% of the variation, was driven primarily by differential usage of G/C- and A/T-ending codons between species. PC2 explained 11.96% of the variation and differentiated the Serinales and Dipodascales orders. The Lipomycetales species at the top left corner, *Dipodascopsis tothii,* is driven by TTG, CTT, CTA, CTG, AGT, TCA, and TCT codons.

*Dipodascopsis tothii* exhibits a long branch length from its relatives and an extremely high GC3 content (91%). The results of this species indicate that it has undergone a unique trajectory in its codon usage that is different from all the rest.

The application of machine learning to genomic data is emerging as a powerful tool for studying yeast traits and evolution (*28, 47*). The pPCA suggested that codon usage can distinguish some, but not all, yeast orders. Machine learning methods, however, can pick up patterns that may be missed in a principal components analysis. To test if codon usage is a distinguishing feature of yeast orders, we constructed a random forest classifier model to classify the order of yeast strains solely on their genome-wide RSCU values (Table S2). The model accurately classified 90.38% of the training data and 93.29% of the test data (30% previously withheld), which indicates that CUB is generally sufficient to differentiate yeast orders (Figure S2). The model was interrogated for the most important variables used for classification using the mean decrease Gini index (Table S3). The most important variable in the model was the codon CTA (leucine), followed by the codons TTG (leucine), AGA (arginine), CTG (leucine), GTA (valine), and CGA (arginine). The relatively high importance of the CTA codon is consistent with the finding that it is a rare case of an A/T ending codon that is generally unpreferred throughout the subphylum. Moreover, this highlights the importance of reassigning the CTG codon in 3 orders. The random forest algorithm could distinguish between genomes belonging to different orders using only the 59 RSCU metrics. The RSCU values, therefore, likely contain significant phylogenetic information.

### Codon Usage Biases Correlated to Genomic Features

Codon usage bias may be shaped by various genomic and ecological factors over the course of evolution. To identify factors shaping codon usage bias of specific codons and overall levels of translational selection, we conducted numerous phylogenetic generalized least squares (PGLS) regressions. These regressions allowed us to account for the fact that our observations share ancestry and are, therefore, a non-random sample.

First, we explored what factors may shape the usage of specific codons by correlating genome-level RSCU values with genomic features, such as other RSCU values, GC content, and tRNA-ome size. We identified 13 codons strongly associated with GC content and tRNA-ome size. Many of these codons were previously identified in the PCA analysis (CTG, AGT, TCT, TTA, CTT; Table S4; Figure S3.). We also tested for codon-to-codon correlations by testing 1,830 pairwise combinations of RSCU (Table S5; Figure S4). After multiple test corrections, 1,785 pairwise combinations were significant at p < 0.05. Of the significant comparisons, 1,740 had the expected correlation between G/C- and A/T-ending codons–they were positively correlated within A/T or G/C comparison and negatively correlated for A/T versus G/C comparisons. There were 23 pairwise comparisons that violated our expectation that G/C- and A/T-ending codons should not exhibit positive correlations in RSCU. Comparisons with high slopes include a positive correlation between TTG (leucine) and CTA (leucine; slope = 1.50) and between AGG (arginine) and CTA (leucine; slope = 0.53). Additionally, 22 comparisons between A/T- or G/C-ending codons were not positively correlated. For example, negative correlations within groups include TCA (serine) and CTA (leucine; slope = -1.20) and AGA (arginine) and CGA (arginine; slope = -0.68). Of the 45 correlations producing results that deviated from the hypothesis that G/C- and A/T-ending codons should be anticorrelated, 43 involved arginine (n=12) or leucine (n=35) codons. This result is consistent with previous studies (*20*) and may be associated with the large number of degenerate codons encoding arginine and leucine, leading to more opportunities for poor codon-tRNA pairing (*48, 49*).

Second, we explored the factors that shape translational selection on codon usage by identifying features that correlate with the S-value (*10*), which is a measure of translational selection (Table S6; Figure S5.) We examined the role of metabolic niche breadth for both carbon and nitrogen using a PGLS analysis. Previous work has shown that intrinsic factors, such as gene composition, drive metabolic niche breadth (*28*). We did not find any association between genome-wide levels of selection on codon usage and carbon (p-value = 0.966) or nitrogen (p-value= 0.579) niche breadth (Table S6). We tested associations between translational selection on codon usage and other genomic features. This includes factors previously associated with selection on codon usage, such as GC3 content, genome size, and tRNA-ome size (*10, 20*). We found positive and significant correlations between the S-value and tRNA-ome (p-value∼ 0) size but not with genome size (p-value = 0.534; Fig 3). Interestingly, both genomic tRNA pool and genome size appeared to serve as lower-bound to high levels of translational selection. Genomes with low levels of translational selection (S-Value < 0.05) were limited to genomes with small genomic tRNA pools. Conversely, high levels of translational selection were found across the genome and tRNA pool size spectrum. The results for synonymous GC3 content were similar to findings in previous studies that suggest the highest levels of translation selection occur at intermediate GC3 content (approximately 50% GC) (*20*).

**Figure 3.**
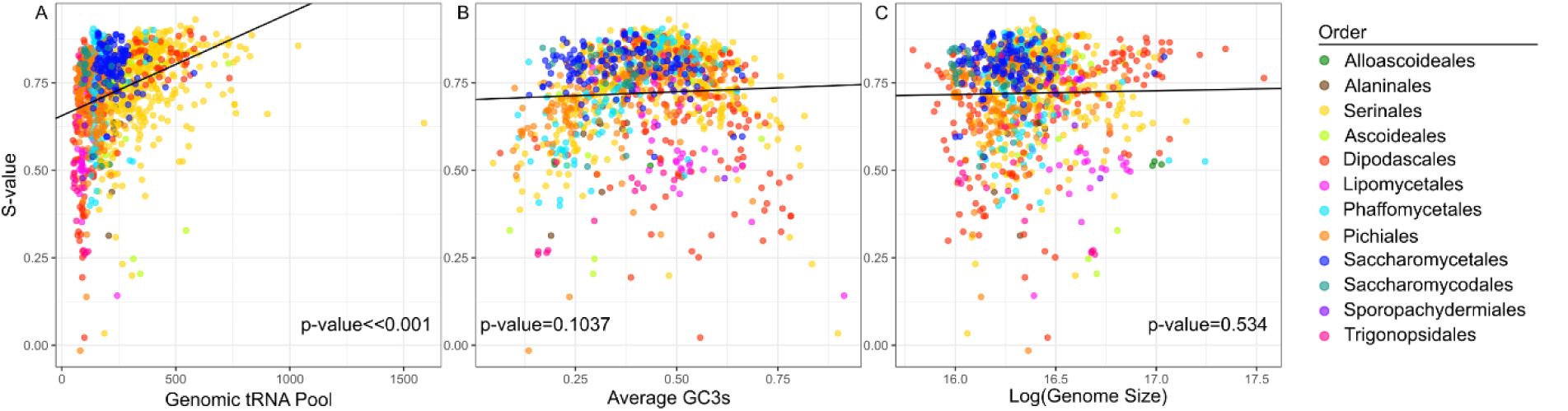
Analysis of 1,154 yeasts reveals a significant association between selection on codon usage bias and tRNA but not with average GC3 content or genome size. (A) PGLS of S-value and tRNA size revealed that there was a significant positive correlation (p-value ∼ 0, slope = 0.00029314). Multiple species in different orders exhibit a wide range of S-values at lower tRNA sizes. Species in orders Dipodascales, Serinales, Saccharomycetales, and Phaffomycetales tended to exhibit higher levels in S-value with increased tRNA size. (B) PGLS of S-value and the average GC content in the third position of the codon that is synonymous were not correlated (p-value = 0.1037, slope = 0.04464). However, visualization suggests a non-linear relationship in which the highest levels of translational selection occur at intermediate GC3 content. C. There was also no association between genome size (log value) and the level of translational selection (p-value = 0.53407, slope= 0.010563).

We also found significant correlations between S-Value and genome assembly metrics (number of contigs, N50), BUSCO metrics (number of BUSCOs, complete, single, fragmented, and missing), and the number of coding sequences (Table S6). We found a positive correlation between S-value and N50 (p-value = 0.000143) and between S-value and the number of BUSCO genes (p-value ∼0), but we found a negative correlation between S-value and the number of contigs (p-value = 0.00193). To further explore the role of all the features identified in pairwise comparisons, we built additive regression models for all possible combinations of tRNA pool size, genome size, N50, number of BUSCO genes, and total number of coding sequences (Table S7). Based on both AIC and BIC criteria, the best-fitting PGLS model included all five variables. This result suggests that either genome assembly quality biases the estimates of selection on codon usage or underlying genomic features that make assemblies more difficult to assemble also impact selection on codon usage. Interestingly, we find that the number of protein-coding sequences is also negatively correlated with the total length of the genome (p-value ∼ 0), the N50 (p-value = 4.40E-12), and the total predicted number of tRNAs (p-value ∼ 0; Table S6). These correlations could indicate that large genomes with a low gene density are harder to assemble and exhibit less translational selection on codon usage. This hypothesis is supported by research that has shown that lower levels of selection associated with low effective population size can lead to larger genomes (*50*). Our results and this model suggest that low levels of selection in this scenario apply to both synonymous and nonsynonymous changes.

### Avoidance of CGN Codons Associated with Arginine tRNA Changes

Certain patterns of RSCU values warranted further examination, such as the observation that the CGN codons exhibited extreme avoidances in multiple clades (Fig. 1). Avoidance of CGN has previously been detected in other fungi, including one yeast mitochondrial genome, *Eremothecium gossypii* (*51*). This extreme bias against CGN led us to investigate the arginine tRNAs. In the results generated from tRNAscan-SE, the arginine tRNAs demonstrated a notable characteristic primarily in the Saccharomycodales. In the *Hanseniaspora* clade, all but three species (*H. singularis*, *H. valbynesis*, and *H. smithiae*) are predicted to be missing the necessary tRNAs to decode CGN codons. However, all the *Hanseniaspora* had predicted tRNAs for AGN codons with an extreme abundance of tRNA-UCU anticodons. The tRNA-UCU anticodons can complementarily base-pair with the codon AGA and AGG through wobble base pairing, which could explain the extreme preference for codon AGA demonstrated in the RSCU heatmap (Fig. 3A) and the low preference of CGN and AGG codons. As noted previously, three *Hanseniaspora* species (*H. singularis*, *H. valbynesis*, and *H. smithiae*) were annotated with a tRNA copy of tRNA-CCG. The presence of this tRNA is particularly interesting as these species are in the fast-evolving lineage (FEL), which has historically experienced significantly higher rates of mutations and gene loss (*52*). The outgroup to the *Hanseniaspora* clade, *Saccharomycodes ludwigii*, is the only species in Saccharomycodales with multiple predicted isotypes to decode CGN codons. The Saccharomycodales mitochondrial genomes were also analyzed for tRNA genes using tRNAscan-SE (Table S8 & Figshare). The only mitochondrial arginine tRNA was tRNA-UCU, which decodes AGA and AGG. This eliminates the possibility that a mitochondrial tRNA is exported to alleviate the nuclear deficiency.

An alternative hypothesis to the degeneration of tRNA genes is that tRNA-scan incorrectly described the CGN decoding tRNAs as pseudogenes. We obtained the sequences for all tRNA isotypes predicted to decode CGN codons regardless of their reported score or predicted isotype (Table S9). All the *Hanseniaspora* have a tRNA with an ACG anticodon (decoding CGT), but the predicted isotype is either lysine, histidine, glycine, or arginine.

Similarly, there are an additional 20 *Hanseniaspora* species that have predicted tRNAs with a CCG anticodon (decoding CGG). These are all predicted to have a histidine or glycine isotype. To determine why tRNA-scan predicted a mismatch between codon and tRNA, we used tRNAviz (*34*) to compare the *Hanseniaspora* sequences to the Saccharomycotina reference.

The alignments of the tRNA sequences revealed positions that were similar and divergent from the consensus sequences (Fig. 4). The *Hanseniaspora* tRNAs with a CCG anticodon diverged from the consensus and *S. ludwigii* at the conserved position 29:41 in the anticodon stem of the tRNA. There is a predicted mismatch here between a G and U base. Previous investigations in *S. cerevisiae* found that tRNA with variants in this location is associated with tRNA decay and maturation dynamics (*53*), and no tRNA modifications have been described at this location (*54*). The *Hanseniaspora* tRNAs with an ACG anticodon diverged from the other sequences at positions 2:71 (mismatch) and 50:64. The mismatch at position 2:71 is located along the acceptor stem, which has been shown to physically interact with the arginyl-tRNA synthetase in *S. cerevisiae* (*55*). This position has also been shown to be a positive identity element for the arginyl-tRNA synthetase in bacteria (*56*). The change at position 50:64 is along the T-arm. The T-arm is not required for tRNA function, suggesting changes in this region may be neutral (*57*).

**Figure 4.**
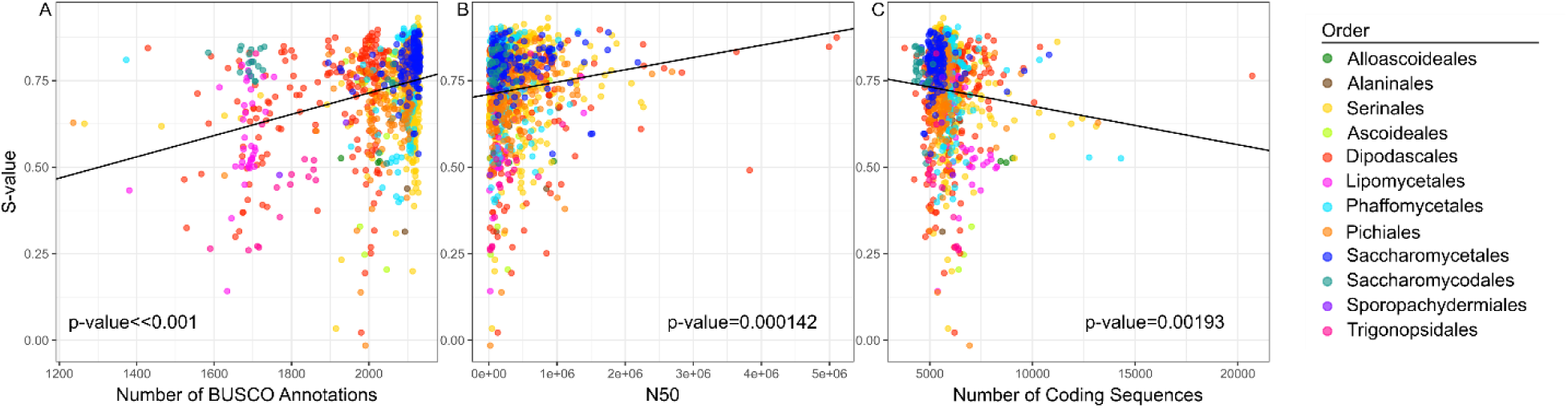
Selection on codon usage is positively correlated with standard measures of genome completeness but not with the total number of protein-coding sequences predicted in a genome. A. Species with larger numbers of BUSCO-annotated genes also had higher levels of translational selection (PGLS, p-value ∼ 0, slope= 0.00030723). B. Species with higher quality genome annotations (measured by N50) also exhibited higher levels of translational selection (PGLS, p-value = 0.00014272, slope = 3.54E-08). C. Unlike the association between the S-value and the number of BUSCO annotations, the total number of coding sequence annotations was negatively correlated with translational selection (PGLS, p-value = 0.0019342, slope = -1.11E-05).

**Figure 5:**
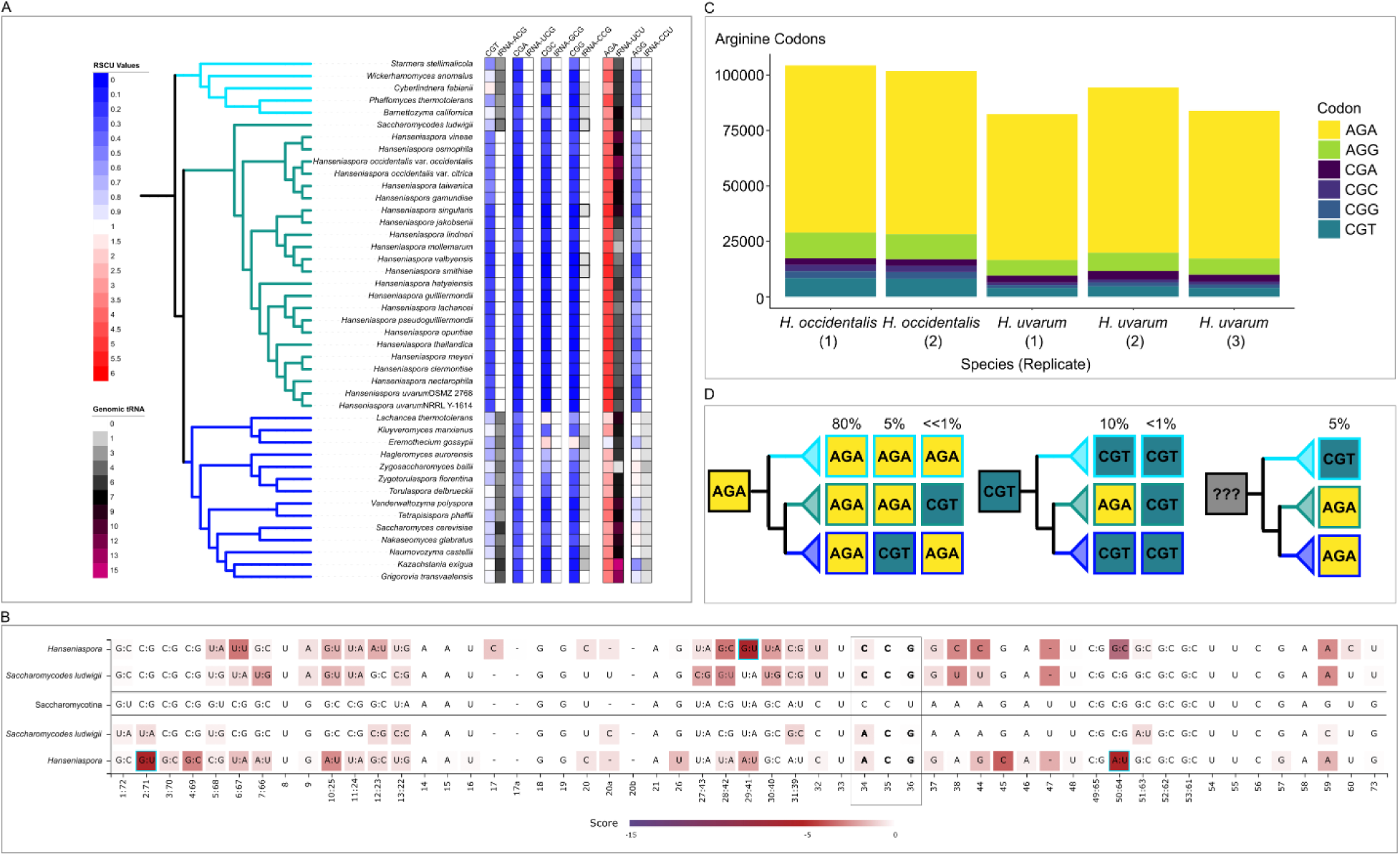
Loss of CGN decoding tRNAs in the *Hanseniaspora* and avoidance, but not complete removal, of CGN codons. (A) The phylogenetic context tRNA genomic content and corresponding RSCU for arginine codons in *Hanseniaspora* species and representative species from Phaffomycetales (top - aqua) and Saccharomycetales (bottom - blue). All four CGN codons are underrepresented in *Hanseniaspora* (RSCU < 1), while the AGA codon is highly overrepresented (RSCU > 1) with the mean *Hanseniaspora* value being 5 out of 6. Three *Hanseniaspora* species have a hypothesized CGG-decoding tRNA, but their tRNA-scan scores are marginal. (B) Comparison of the annotated CCG and ACG anti-codon-containing tRNAs and pseudo-tRNAs compared to the rest of the Saccharomycotina. The dark red locations indicate positions that are divergent from the reference sequence shown in the middle. (C) The total number of arginine codons that were identified in five transcriptomes in two *Hanseniaspora* species. The majority of identified codons were AGN, but there were still CGN codons detected in the mRNA expression data. (D) The evolutionary history of conserved arginine positions (80% across Saccharomycotina) in orthologous genes that are also conserved within the Phaffomycetales (top), Saccharomycodales (middle), and Saccharomycetales (bottom). CGN codons are rare overall at conserved positions. At positions where the ancestral codon was likely CGT, the vast majority of these positions evolved to AGA (134/137). Overall, there were only 4 sites out of 1,404 that retained CGT across the Saccharomycodales.

Additionally, many *Hanseniaspora* genomes have lost the tRNA adenosine deaminase (*TAD*) genes, which include homologs that have been shown to modify the wobble position arginine tRNAs with an ACG anticodon (decoding CGT codons) (*58*) (Table S10). The enzyme encoded by *TAD1* is found in 9 of the 24 (38%) Saccharomycodales species, a fraction that is significantly lower than across the rest of the yeast subphylum (1065/1130 or 94%, Chi-squared with Yates’ continuity correction p-value < 2.2e-16) or when compared to the Saccharomycetales (134/135 or 99%, Chi-squared with Yates’ continuity correction p-value < 2.2e-16). The enzymes encoded by *TAD2* and *TAD3* form a heterodimer (*59*). In the Saccharomycodales, 21 out of 24 yeast (88%) have both components, which is comparable to the rest of the subphylum (963/1130 or 85%). This tRNA modification enzyme complex modifies the wobble position (A) to inosine (I) to allow for better wobble pairing (*25*). While additional evidence would be required to demonstrate that the *Hanseniaspora TAD* genes modify arginine tRNAs, this loss further narrows down possible explanations for how these species decode CGN codons.

While the CGN codons are rare in the *Hanseniaspora* species without a predicted tRNA (mean RSCU across all CGN of 0.177), they are not completely absent. If the tRNAs were completely non-functional, we would expect that genes containing CGN codons would be untranslatable, leading to their extinction or elimination of CGN codons. To determine if transcripts containing CGN codons are expressed, we conducted RNA sequencing of *H. occidentalis* var. *occidentalis* and *H. uvarum* cultured in a rich glucose medium (Fishare & GenBank). We then counted the total number of arginine codons in the expressed genes. The majority of arginine codons in the transcripts were AGA (mean 76% across samples). The CGN codons only comprised 14% (mean across samples) of the total arginine codons. At the level of transcripts, 37% of genes expressed contained no CGN codons. Of the remaining transcripts that did contain CGN codons, the median number of CGN codons was 4, and the mean was 7.7. Most of the CGN codons are concentrated into a few transcripts, with 1% (258 of 22498) of the transcripts containing more than 20 CGN codons. We investigated the 20 transcripts with the most CGN codons by BLASTing (tblastx) against the non-redundant protein database database (Table S11). Twelve transcripts had high similarity to Saccharomycodales genomes but not protein sequences, suggesting that they are either non-coding transcripts or previously unannotated protein sequences. The remaining eight sequences had partial matches to previously reported Saccharomycodales protein-coding genes. The sequence with the highest percent identity to a known gene matched the *FLO1* gene, which encodes a flocculation protein from *H. uvarum*. The relatively high number of CGN codons in this gene (62 codons) may be due to the previously characterized extended tandem repeats in this gene (*60*). The low average percent identity (55%) of these 20 transcripts to known genes suggests they are not likely to be complete translated mRNA sequences. As a comparison, we also analyzed 20 randomly chosen transcripts with no CGN codons. In this case, 18 of the 20 translations had high similarity (average of 91%) to previously characterized proteins in the *Hanseniaspora*. Only two sequences showed no significant similarity in the BLAST database. This suggests that the CGN-containing transcripts from our sequencing experiment are not protein-coding mRNA transcripts.

Our analysis indicated that CGN codons are also generally avoided, albeit to a lesser degree, in Phaffomycetales and Saccharomycetales, the two orders most closely related to the Saccharomycodales. To examine if the Saccharomycodales are evolving away from CGN codons compared to their relatives, we compared conserved arginine codon positions. These positions were identified in the 1,403 conserved orthologs used to determine the Saccharomycotina phylogeny. Positions were required to be arginine in 80% of all the sequences and the same codon 80% of the time within each order. This allowed us to examine 1,404 conserved arginine positions in 1,400 orthologs (Figshare Repository). We then determined the most parsimonious ancestral codon across the orders. In ∼85% of the positions, the inferred ancestor was AGA. In only one conserved position did we infer a change from AGA to CGT in the Saccharomycodales. In positions where CGT was the ancestral codon, the CGT codon was retained in only 3 positions, while 134 transitioned to AGA. We also conducted this analysis with positions that were conserved at a lower threshold (60%) and found similar results (Table S12). This analysis suggests that, in conserved arginine positions, the Saccharomycodales have repeatedly switched codons from CGN to AGN. The change from CGN to AGN requires two base pair mutations in 6 of the 8 possible ways to go from CGN to AGN.

Collectively, our analysis found that the extreme avoidance of CGN codons in *Hanseniaspora* was likely associated with the accumulation of mutations in the CGN-decoding tRNAs. The transcriptomics data suggests that transcripts containing CGN codons are rare, with over a third of transcripts containing zero CGN codons. This phenomenon may be associated with the loss of DNA repair and cell cycle genes previously observed in this group (*52*). This scenario resembles the hypothesized situation leading to the CTG codon reassignment in three orders. In the “tRNA loss driven codon reassignment” model of CTG codon reassignment, loss of function mutations in tRNAs is the driving factor in codon reassignment. The CGN tRNAs appear to have accumulated several mutations, making them unrecognizable to tRNAscan-SE (*61*). An alternative hypothesis is that codon reassignment is driven by “codon capture,” in which a codon is driven to near extinction before changes in tRNAs (*62*). Additional work is needed to test the expression or modification of the CGN tRNAs in *Hanseniaspora*.

## Conclusions

The Saccharomycotina exhibit vast diversity in their codon usage and genomic tRNA content. Each order has evolved distinct codon usage patterns–including codon reassignments– that are sufficiently divergent to classify yeasts into their order using RSCU alone. Many forces shape the codon usage of Saccharomycotina, including mutational bias, genomic tRNA pool, and overall genome content. Similar to previous studies (*18, 20, 27*), we find that the genomic tRNA pool serves as a lower bound for the amount of translational selection acting on codon usage–small pools can exhibit a range of S-values. In contrast, large pools exhibit mostly high S-values. We also find that the highest levels of translational selection occur at an intermediate GC content of the third codon position. Interestingly, we find that the N50, BUSCO number, and translational selection are all positively correlated with each other and negatively correlated with the total number of predicted protein-coding sequences.

Unlike previous studies, we identified an association between genome assembly and architecture and our measure of translational selection. Our additive model found that 5 variables (tRNA pool size, genome size, N50, number of BUSCO genes, and total number of protein-coding sequences) explained the most variation in S-value. The role of genome assembly features could be technical or biological. Lower-quality genomes may be missing genes and are more likely to be misannotated. This could lead to unintentional bias in the codon evaluation due to missing genes or tRNAs (*63*). Conversely, biological genome features, like high GC content (*64*), repetitive regions, presence of introns, heterozygosity, and genome size (*65*) can result in lower-quality genome assemblies. Many of these features, like GC content and genome size, have previously been found to be associated with codon usage bias (*10, 20, 26*). Therefore, the improved model fit associated with adding features like N50 and BUSCO may be associated with genome features we did not capture in our model.

Our analysis also uncovered an extreme avoidance of the CGN arginine codons in the Saccharomycodales. This was associated with a widespread predicted loss of function in the *Hanseniaspora* tRNAs, which decode CGN codons. Despite this observation, RNA-sequencing data identified several transcripts rich in CGN codons. However, whether these transcripts result in amino acids or the CGN tRNAs are expressed within the cell remains to be seen. Overall, the tRNAs that decode CGN codons have accumulated multiple mutations that may impact their function. The *Hanseniaspora* have also generally lost the Tad1 enzyme, and three have lost the ability to form the Tad2/Tad3 heterodimer—these enzymes are involved in modifying tRNAs to increase wobble base pairing (*58, 66*)The evolution away from CGN codons, the accumulation of mutations in the CGN-decoding tRNAs, and the loss of the *TAD1* genes all support the hypothesis that many *Hanseniaspora* species have a significantly impaired ability to decode CGN codons.

Our analysis of codon usage bias in the Saccharomycotina revealed diverse codon usage biases, widespread selection on codon usage, and an extreme avoidance of CGN codons in an order that has potentially lost the tRNAs to decode CGN codons. Given the diversity in codon usage, the subphylum will likely be critical in answering outstanding questions in the field of codon usage. The *Hanseniaspora* may allow us to observe codon reassignment in action. The various species with incredibly large and very small numbers of tRNA genes may help us answer questions about the role of tRNA copy number and sequence variation in regulation. Finally, as we learn more about the ecology of these yeasts, we may be able to identify life history traits that impact selection on codon usage.

## Author Contributions

BZ conducted computational and statistical analyses, managed data, prepared figures, and co-wrote the manuscript with ALL.

LD conducted tRNA modification enzyme analysis.

KJF generated *Hanseniaspora* mRNA sequencing data.

DAO, XXS, XZ, JFW, MCH, MZ, CTH, and AR provided computational support and reagents.

ALL designed and implemented computational analyses, managed data, prepared figures, co-wrote the manuscript, and supervised the project.

All authors provided comments and input and approved the manuscript.

## Acknowledgements

Computational analyses were run in the UNC Charlotte high performance computing cluster in Charlotte North Carolina. X.-X.S. was supported by the NSF for Distinguished Young Scholars of Zhejiang Province (LR23C140001), the Fundamental Research Funds for the Central Universities (226-2023-00021), and the key research project of Zhejiang Lab (2021PE0AC04). This work was supported by the NSF (grants DEB-2110403 to C.T.H. and DEB-2110404 to A.R.). Research in the Hittinger Lab is also supported by the USDA National Institute of Food and Agriculture (Hatch Projects 1020204 and 7005101), in part by the DOE Great Lakes Bioenergy Research Center [DOE BER Office of Science DE–SC0018409, and an H.I. Romnes Faculty Fellowship (Office of the Vice Chancellor for Research and Graduate Education with funding from the Wisconsin Alumni Research Foundation)]. Research in the Rokas lab is also supported by the NIH/National Institute of Allergy and Infectious Diseases (R01 AI153356), and the Burroughs Wellcome Fund.

## Data and materials availability

The Y1000+ data can be obtained from the project website (https://y1000plust.org). The Figshare repository contains the raw random forest model data, the assembled transcriptomes from the *Hanseiaspora*, the RSCU for all coding sequences in the subphylum, the conserved argninine analysis, and the mitochondrial tRNA analysis.

The Figshare repository and genbank accession numbers for the *Hanseniaspora* transcriptome data will be provided upon publication.

## Works Cited

1. T. Ikemura, Codon usage and tRNA content in unicellular and multicellular organisms. Mol Biol Evol 2, 13–34 (1985).

2. J. B. Plotkin, G. Kudla, Synonymous but not the same: the causes and consequences of codon bias. Nat Rev Genet 12, 32–42 (2011).

3. R. Grantham, Viral, prokaryote and eukaryote genes contrasted by mRNA sequence indexes. FEBS Lett 95, 1–11 (1978).

4. P. M. Sharp, T. M. Tuohy, K. R. Mosurski, Codon usage in yeast: cluster analysis clearly differentiates highly and lowly expressed genes. Nucleic Acids Res 14, 5125–5143 (1986).

5. A. Radhakrishnan et al., The DEAD-Box Protein Dhh1p Couples mRNA Decay and Translation by Monitoring Codon Optimality. Cell 167, 122–132 e129 (2016).

6. A. Coghlan, K. H. Wolfe, Relationship of codon bias to mRNA concentration and protein length in Saccharomyces cerevisiae. Yeast 16, 1131–1145 (2000).

7. M. Zhou, T. Wang, J. Fu, G. Xiao, Y. Liu, Nonoptimal codon usage influences protein structure in intrinsically disordered regions. Mol Microbiol 97, 974–987 (2015).

8. F. Zhao et al., Genome-wide role of codon usage on transcription and identification of potential regulators. Proc Natl Acad Sci U S A 118, (2021).

9. C. H. Yu et al., Codon Usage Influences the Local Rate of Translation Elongation to Regulate Co-translational Protein Folding. Mol Cell 59, 744–754 (2015).

10. M. dos Reis, R. Savva, L. Wernisch, Solving the riddle of codon usage preferences: a test for translational selection. Nucleic Acids Res 32, 5036–5044 (2004).

11. Y. Lesecque, D. Mouchiroud, L. Duret, GC-biased gene conversion in yeast is specifically associated with crossovers: molecular mechanisms and evolutionary significance. Mol Biol Evol 30, 1409–1419 (2013).

12. H. Liu et al., Tetrad analysis in plants and fungi finds large differences in gene conversion rates but no GC bias. Nat Ecol Evol 2, 164–173 (2018).

13. Y. O. Zhu, M. L. Siegal, D. W. Hall, D. A. Petrov, Precise estimates of mutation rate and spectrum in yeast. Proc Natl Acad Sci U S A 111, E2310–2318 (2014).

14. M. Lynch et al., A genome-wide view of the spectrum of spontaneous mutations in yeast. Proc Natl Acad Sci U S A 105, 9272–9277 (2008).

15. Y. I. Pavlov, C. S. Newlon, T. A. Kunkel, Yeast origins establish a strand bias for replicational mutagenesis. Mol Cell 10, 207–213 (2002).

16. N. Galtier et al., Codon Usage Bias in Animals: Disentangling the Effects of Natural Selection, Effective Population Size, and GC-Biased Gene Conversion. Mol Biol Evol 35, 1092–1103 (2018).

17. G. Boel et al., Codon influence on protein expression in E. coli correlates with mRNA levels. Nature 529, 358–363 (2016).

18. C. Landerer, A. Cope, R. Zaretzki, M. A. Gilchrist, AnaCoDa: analyzing codon data with Bayesian mixture models. Bioinformatics 34, 2496–2498 (2018).

19. M. A. Gilchrist, W. C. Chen, P. Shah, C. L. Landerer, R. Zaretzki, Estimating Gene Expression and Codon-Specific Translational Efficiencies, Mutation Biases, and Selection Coefficients from Genomic Data Alone. Genome Biol Evol 7, 1559–1579 (2015).

20. A. L. LaBella, D. A. Opulente, J. L. Steenwyk, C. T. Hittinger, A. Rokas, Variation and selection on codon usage bias across an entire subphylum. PLoS Genet 15, e1008304 (2019).

21. R. L. Nalabothu et al., Codon optimization improves the prediction of xylose metabolism from gene content in budding yeasts. Mol Biol Evol, (2023).

22. A. L. LaBella, D. A. Opulente, J. L. Steenwyk, C. T. Hittinger, A. Rokas, Signatures of optimal codon usage in metabolic genes inform budding yeast ecology. PLoS Biol 19, e3001185 (2021).

23. S. Muhlhausen, P. Findeisen, U. Plessmann, H. Urlaub, M. Kollmar, A novel nuclear genetic code alteration in yeasts and the evolution of codon reassignment in eukaryotes. Genome Res 26, 945–955 (2016).

24. R. Riley et al., Comparative genomics of biotechnologically important yeasts. Proc Natl Acad Sci U S A 113, 9882–9887 (2016).

25. M. Wada, K. Ito, The CGA codon decoding through tRNA(Arg) (ICG) supply governed by Tad2/Tad3 in Saccharomyces cerevisiae. FEBS J 290, 3480–3489 (2023).

26. A. L. Cope, P. Shah, Intragenomic variation in non-adaptive nucleotide biases causes underestimation of selection on synonymous codon usage. PLoS Genet 18, e1010256 (2022).

27. R. Wint, A. Salamov, I. V. Grigoriev, Kingdom-Wide Analysis of Fungal Protein-Coding and tRNA Genes Reveals Conserved Patterns of Adaptive Evolution. Mol Biol Evol 39, (2022).

28. D. A. Opulente et al., Genomic factors shape carbon and nitrogen metabolic niche breadth across Saccharomycotina yeasts. Science 384, eadj4503 (2024).

29. P. Rice, I. Longden, A. Bleasby, EMBOSS: the European Molecular Biology Open Software Suite. Trends Genet 16, 276–277 (2000).

30. P. P. Chan, B. Y. Lin, A. J. Mak, T. M. Lowe, tRNAscan-SE 2.0: improved detection and functional classification of transfer RNA genes. Nucleic Acids Res 49, 9077–9096 (2021).

31. M. A. Santos, M. F. Tuite, The CUG codon is decoded in vivo as serine and not leucine in Candida albicans. Nucleic Acids Res 23, 1481–1486 (1995).

32. T. Krassowski et al., Evolutionary instability of CUG-Leu in the genetic code of budding yeasts. Nat Commun 9, 1887 (2018).

33. J. F. Wolters, A. L. LaBella, D. A. Opulente, A. Rokas, C. T. Hittinger, Mitochondrial genome diversity across the subphylum Saccharomycotina. Front Microbiol 14, 1268944 (2023).

34. B. Y. Lin, P. P. Chan, T. M. Lowe, tRNAviz: explore and visualize tRNA sequence features. Nucleic Acids Res 47, W542–W547 (2019).

35. M. Kanehisa, S. Goto, KEGG: kyoto encyclopedia of genes and genomes. Nucleic Acids Res 28, 27–30 (2000).

36. Z. Gu, R. Eils, M. Schlesner, Complex heatmaps reveal patterns and correlations in multidimensional genomic data. Bioinformatics 32, 2847–2849 (2016).

37. L. J. Revell, phytools 2.0: an updated R ecosystem for phylogenetic comparative methods (and other things). PeerJ 12, e16505 (2024).

38. D. Orme et al., The caper package: comparative analysis of phylogenetics and evolution in R. R package version 5, 1–36 (2013).

39. E. Paradis, K. Schliep, ape 5.0: an environment for modern phylogenetics and evolutionary analyses in R. Bioinformatics 35, 526–528 (2019).

40. A. Liaw, M. Wiener, Classification and regression by randomForest. R news 2, 18–22 (2002).

41. S. Chen, Ultrafast one-pass FASTQ data preprocessing, quality control, and deduplication using fastp. Imeta 2, e107 (2023).

42. D. Kim, J. M. Paggi, C. Park, C. Bennett, S. L. Salzberg, Graph-based genome alignment and genotyping with HISAT2 and HISAT-genotype. Nat Biotechnol 37, 907–915 (2019).

43. A. Shumate, B. Wong, G. Pertea, M. Pertea, Improved transcriptome assembly using a hybrid of long and short reads with StringTie. PLoS Comput Biol 18, e1009730 (2022).

44. T. Madden, The BLAST sequence analysis tool. The NCBI handbook, (2003).

45. K. Katoh, K. Misawa, K. Kuma, T. Miyata, MAFFT: a novel method for rapid multiple sequence alignment based on fast Fourier transform. Nucleic Acids Res 30, 3059–3066 (2002).

46. C. Marck, H. Grosjean, tRNomics: analysis of tRNA genes from 50 genomes of Eukarya, Archaea, and Bacteria reveals anticodon-sparing strategies and domain-specific features. Rna 8, 1189–1232 (2002).

47. M. C. Harrison et al., Machine learning enables identification of an alternative yeast galactose utilization pathway. Proc Natl Acad Sci U S A 121, e2315314121 (2024).

48. L. Duret, D. Mouchiroud, Expression pattern and, surprisingly, gene length shape codon usage in Caenorhabditis, Drosophila, and Arabidopsis. Proc Natl Acad Sci U S A 96, 4482–4487 (1999).

49. G. A. McVean, J. Vieira, Inferring parameters of mutation, selection and demography from patterns of synonymous site evolution in Drosophila. Genetics 157, 245–257 (2001).

50. D. A. Petrov, Mutational equilibrium model of genome size evolution. Theor Popul Biol 61, 531–544 (2002).

51. M. Carullo, X. Xia, An extensive study of mutation and selection on the wobble nucleotide in tRNA anticodons in fungal mitochondrial genomes. J Mol Evol 66, 484–493 (2008).

52. J. L. Steenwyk et al., Extensive loss of cell-cycle and DNA repair genes in an ancient lineage of bipolar budding yeasts. PLoS Biol 17, e3000255 (2019).

53. M. J. Payea, A. C. Hauke, T. De Zoysa, E. M. Phizicky, Mutations in the anticodon stem of tRNA cause accumulation and Met22-dependent decay of pre-tRNA in yeast. Rna 26, 29–43 (2020).

54. T. Suzuki, The expanding world of tRNA modifications and their disease relevance. Nat Rev Mol Cell Biol 22, 375–392 (2021).

55. B. Delagoutte, D. Moras, J. Cavarelli, tRNA aminoacylation by arginyl-tRNA synthetase: induced conformations during substrates binding. EMBO J 19, 5599–5610 (2000).

56. R. Giege, G. Eriani, The tRNA identity landscape for aminoacylation and beyond. Nucleic Acids Res 51, 1528–1570 (2023).

57. N. Krahn, J. T. Fischer, D. Soll, Naturally Occurring tRNAs With Non-canonical Structures. Front Microbiol 11, 596914 (2020).

58. J. Wolf, A. P. Gerber, W. Keller, tadA, an essential tRNA-specific adenosine deaminase from Escherichia coli. EMBO J 21, 3841–3851 (2002).

59. G. S. Dance et al., Identification of the yeast cytidine deaminase CDD1 as an orphan C-->U RNA editase. Nucleic Acids Res 29, 1772–1780 (2001).

60. F. Bidard, M. Bony, B. Blondin, S. Dequin, P. Barre, The Saccharomyces cerevisiae FLO1 flocculation gene encodes for a cell surface protein. Yeast 11, 809–822 (1995).

61. M. Kollmar, S. Muhlhausen, Nuclear codon reassignments in the genomics era and mechanisms behind their evolution. Bioessays 39, (2017).

62. S. Osawa, T. H. Jukes, Codon reassignment (codon capture) in evolution. J Mol Evol 28, 271–278 (1989).

63. A. Whibley, J. L. Kelley, S. R. Narum, The changing face of genome assemblies: Guidance on achieving high-quality reference genomes. Mol Ecol Resour 21, 641–652 (2021).

64. Y. C. Chen, T. Liu, C. H. Yu, T. Y. Chiang, C. C. Hwang, Effects of GC bias in next-generation-sequencing data on de novo genome assembly. PLoS One 8, e62856 (2013).

65. A. A. Jauhal, R. D. Newcomb, Assessing genome assembly quality prior to downstream analysis: N50 versus BUSCO. Mol Ecol Resour 21, 1416–1421 (2021).

66. E. Delannoy et al., Arabidopsis tRNA adenosine deaminase arginine edits the wobble nucleotide of chloroplast tRNAArg(ACG) and is essential for efficient chloroplast translation. Plant Cell 21, 2058–2071 (2009).

