## Supplemental Figures for "Genomic factors shaping codon usage across the Saccharomycotina subphylum"

Author Contributions:

BZ conducted computational and statistical analyses, managed data, prepared figures, and co-wrote the manuscript with ALL.

LD conducted tRNA modification enzyme analysis.

KJF generated *Hanseniaspora* mRNA sequencing data.

DAO, XXS, XZ, JFW, MCH, MZ, CTH, and AR provided computational support and reagents.

ALL designed and implemented computational analyses, managed data, prepared figures, co-wrote the manuscript, and supervised the project.

All authors provided comments and input and approved the manuscript.

### Supplemental Materials

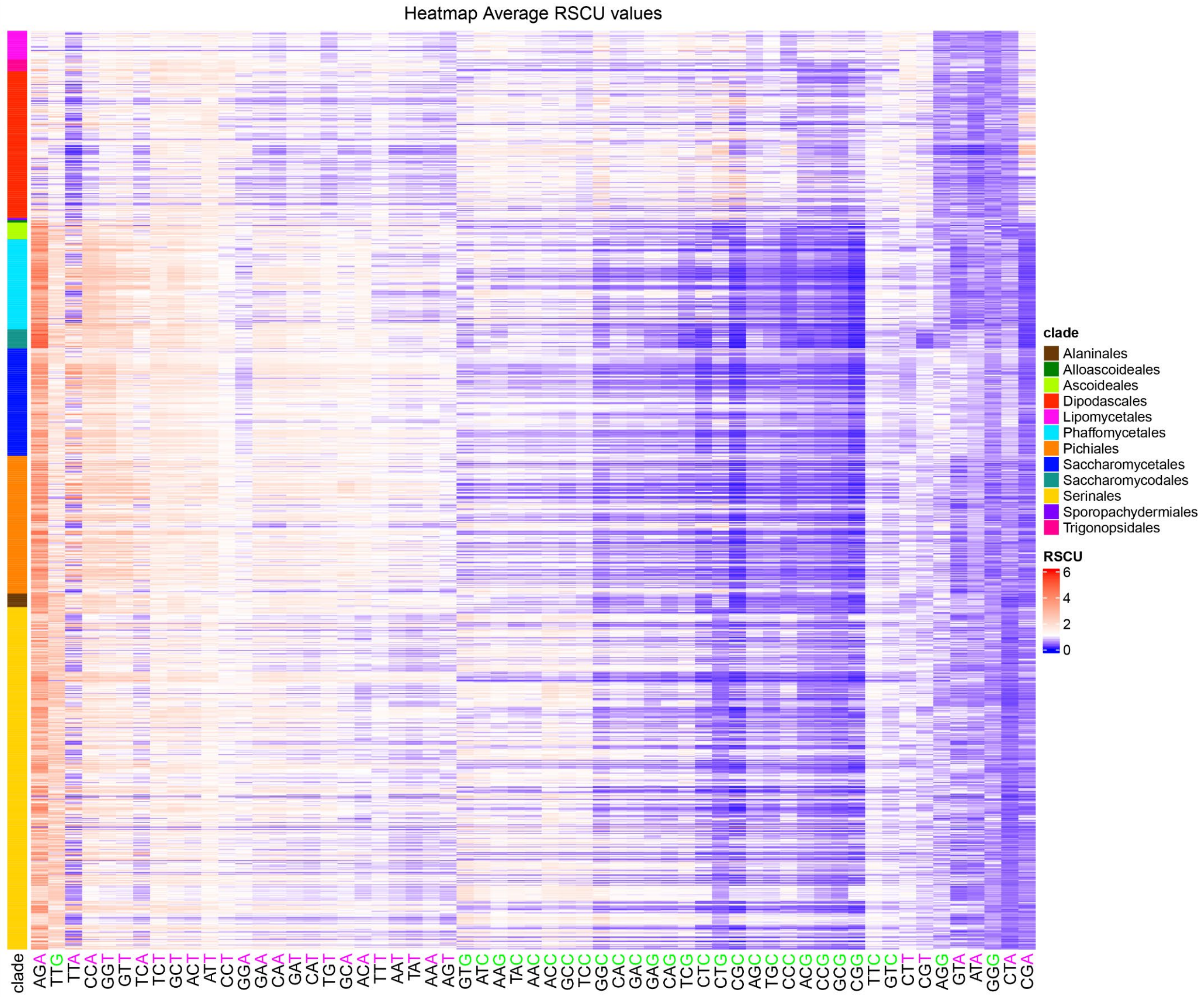


**Supplemental Figure 1: Variation in Relative Synonymous Codon Usage (RSCU) across the yeast subphylum.** Codons in blue are non-preferred, while codons in red are preferred. The third codon position is colored by GC- (green) or AT- (purple) ending codons. The x-axis is clustered based on a hierarchical clustering of RSCU values. The y-axis is sorted by Saccharomycotina order. Generally, GC- and AT-ending codons are clustered together. The exceptions are the general preference for the TTG codon and avoidance of CTT, CGT, GTA, ATA, CTA, and CGA codons.

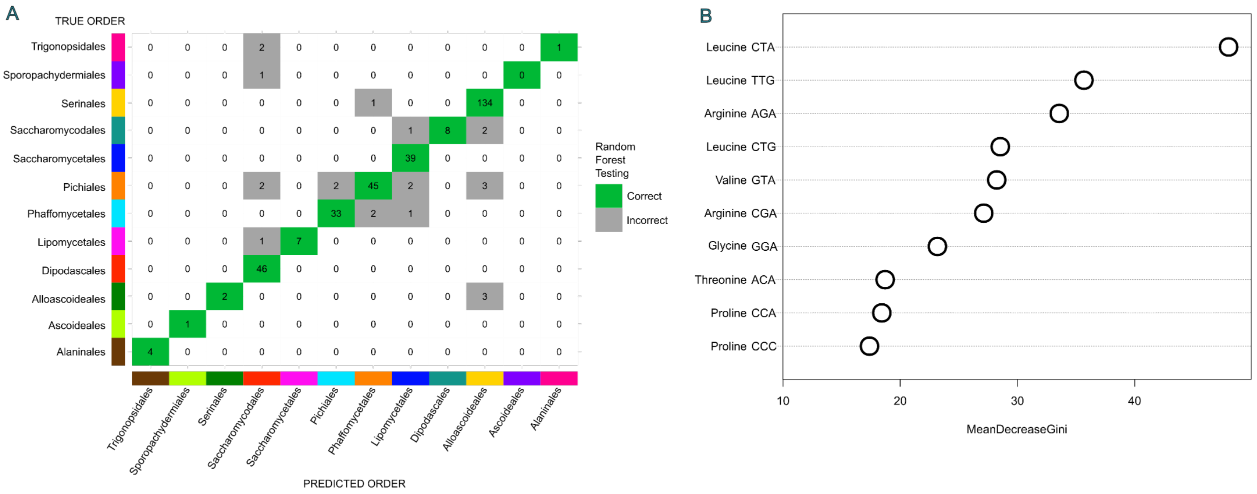


**Supplemental Figure 2:** Random forest classifier sucesfully classifies the majority of the withheld strains using genome-wide RSCU values. A) The confusion matrix of the testing species. The largest number of misclassified yeasts were assigned to the Alloacoideales. B) The relative importance of the codons with the highest importance.

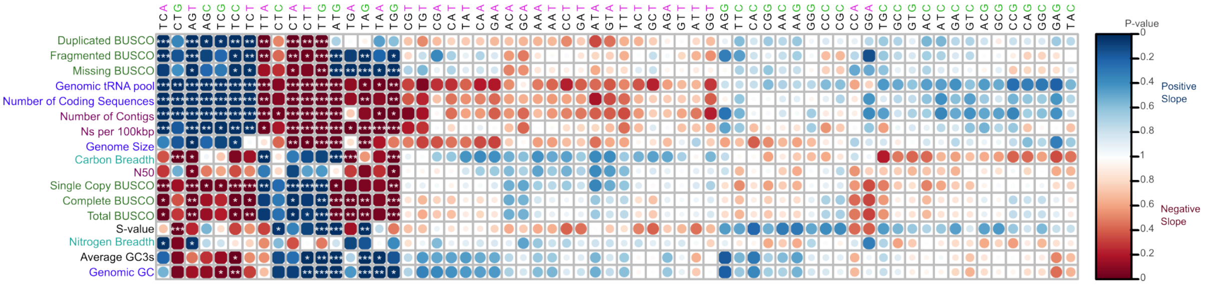


**Supplemental Figure 3:** Correlation between RSCU of all codons and yeast features. Raw p-values are shown on the scale, with positive correlations shown in blue and negative correlations shown in red. More significant correlations are darker with larger circles. Significant raw p-values are shown with stars at the levels 0.05, 0.01, and 0.001.

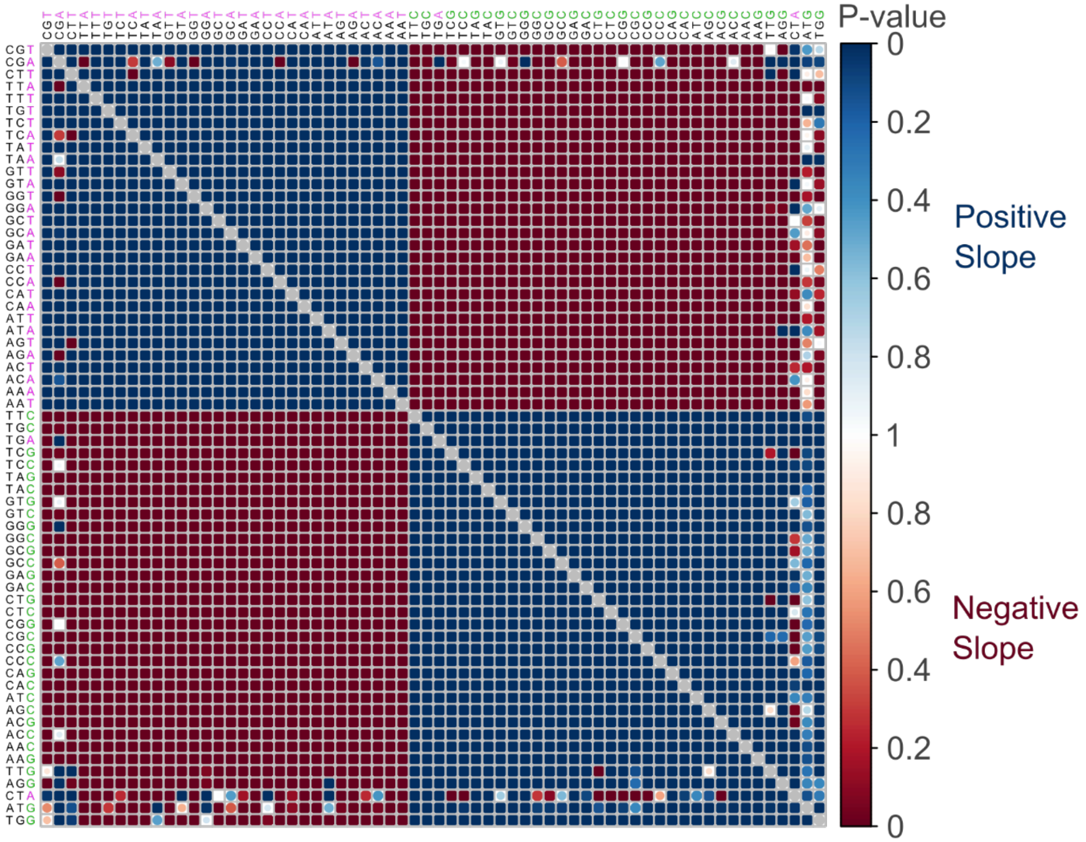


**Supplemental Figure 4:** Correlation between RSCU of all possible codons. Raw p-values are shown on the scale, with positive correlations shown in blue and negative correlations shown in red. More significant correlations are darker with larger circles. Generally, AT- and GC-ending codons are correlated within groups and anti-correlated across groups. The coding sequence with the most unexpected correlations is CTA (leucine).

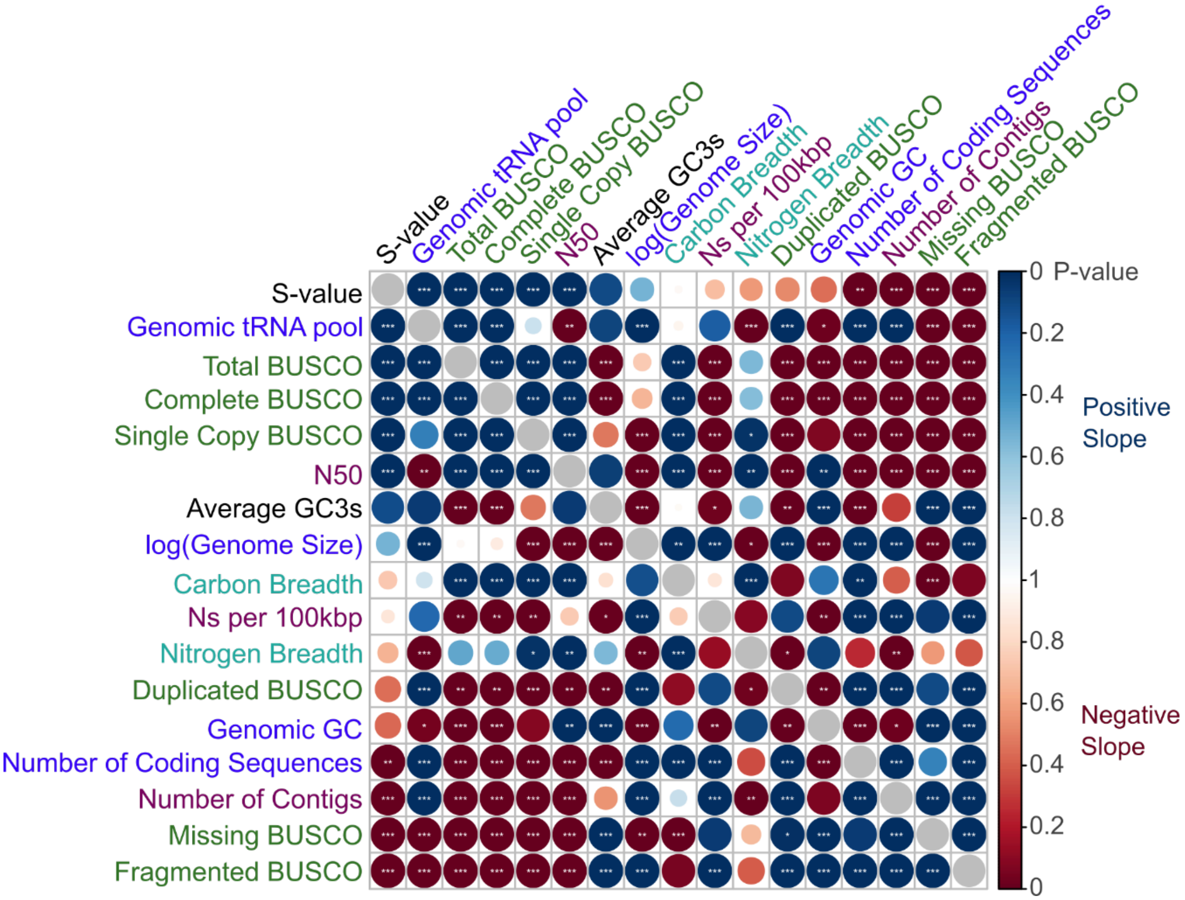


**Supplemental Figure 5:** Correlation between features of the 1,154 yeasts. Raw p-values are shown on the scale, with positive correlations shown in blue and negative correlations shown in red. More significant correlations are darker with larger circles. Significant raw p-values are shown with stars at the levels 0.05, 0.01, and 0.001. S-value has significant positive correlations with genomic tRNA pool, total BUSCO count, complete BUSCO count, single copy BUSCO count, and N50. S-value has significant negative correlations with fragmented BUSCO count, missing BUSCO count, total number of contigs, and number of coding sequences.

**Table S1: Table of yeast genomes and features**. This table includes the origin of the yeast genomes, updated species names, and the examined genomic features.

**Table S2: RSCU of all codons across yeasts.** For each of the 1,154 yeast genome annotations, we report the mean RSCU across all coding sequences for all 61 codons.

**Table S3: Variable importance of random forest algorithm**: Importance values extracted from the random forest algorithm, which classified yeasts into orders based on RSCU. The higher the importance value the more informative it was in building the random forest.

**Table S4: Results of feature versus RSCU comparisons**: The RSCU for each codon was compared to yeast features. The p-value and slope are reported for each comparison.

**Table S5: Results of pairwise RSCU comparisons**: The RSCU for each codon was compared to every other codon using a PGLS. The p-value and slope are reported for each comparison.

**Table S6: Results of pairwise comparisons between S-value and other features**: The S-value was compared to other features using a PGLS. The p-value and slope are reported for each comparison.

**Table S7: Additive PGLS models**. We conducted all possible iterations of additive models to explain S-value based on genomic and assembly metrics. The models' AIC and BIC values are reported and used to select the most informative model accounting for the additional parameters.

**Table S8: Analysis of mitochondrial tRNAs**: This table includes the output from tRNAscan-SE from the available mitochondrial genomes of the *Hanseniaspora*.

**Table S9: All tRNAs with a predicted CGN anticodon within the *Hanseniaspora*.** tRNAscan-SE predicts the majority of tRNAs containing a CGN anticodon to be non-functional. This can be seen in the isotype prediction.

**Table S10: Presence of *TAD*-encoded tRNA modification enzymes in the *Hanseniaspora* and relatives.**

**Table S11: *Hanseniaspora* transcriptomics BLAST results.** The highest BLASTx and BLASTn hits from transcripts with many and with no CGN codons.

**Table S12: Conservation of CGN codons in the *Hanseniaspora*.** Rate of CGN codons in conserved arginine positions reported for the *Hanseniaspora* and their relatives.

947

948
